# Neuropathy-related mutations alter the membrane binding properties of the human myelin protein P0 cytoplasmic tail

**DOI:** 10.1101/535013

**Authors:** Arne Raasakka, Salla Ruskamo, Robert Barker, Oda C. Krokengen, Guro H. Vatne, Cecilie K. Kristiansen, Erik I. Hallin, Maximilian W.A. Skoda, Ulrich Bergmann, Hanna Wacklin-Knecht, Nykola C. Jones, Søren Vrønning Hoffmann, Petri Kursula

**Affiliations:** Department of Biomedicine, University of Bergen, Bergen, Norway; Faculty of Biochemistry and Molecular Medicine, University of Oulu, Oulu, Finland; Biocenter Oulu, University of Oulu, Oulu, Finland; School of Physical Sciences, University of Kent, Kent, United Kingdom; ISIS Neutron and Muon Source, Science & Technology Facilities Council, Rutherford Appleton Laboratory, OX11 OQX Didcot, United Kingdom; Division of Physical Chemistry, Department of Chemistry, Lund University & European Spallation Source ERIC, Lund, Sweden; ISA, Department of Physics and Astronomy, Aarhus University, Ny Munkegade 120, 8000 Aarhus C, Denmark

**Keywords:** Myelin protein zero, membrane binding, peripheral neuropathy, CMT, DSS, disease mutation, gain of function

## Abstract

Schwann cells myelinate selected axons in the peripheral nervous system (PNS) and contribute to fast saltatory conduction *via* the formation of compact myelin, in which water is excluded from between tightly adhered lipid bilayers. Peripheral neuropathies, such as Charcot-Marie-Tooth disease (CMT) and Dejerine-Sottas syndrome (DSS), are incurable demyelinating conditions that result in pain, decrease in muscle mass, and functional impairment. Many Schwann cell proteins, which are directly involved in the stability of compact myelin or its development, are subject to mutations linked to these neuropathies. The most abundant PNS myelin protein is protein zero (P0); point mutations in this transmembrane protein cause CMT subtype 1B and DSS. P0 tethers apposing lipid bilayers together through its extracellular immunoglobulin-like domain. Additionally, P0 contains a cytoplasmic tail (P0ct), which is membrane-associated and contributes to the physical properties of the lipid membrane. Six CMT- and DSS-associated missense mutations have been reported in P0ct. We generated recombinant disease mutant variants of P0ct and characterized them using biophysical methods. Compared to wild-type P0ct, some mutants have negligible differences in function and folding, while others highlight functionally important amino acids within P0ct. For example, the D224Y variant of P0ct induced tight membrane multilayer stacking. Our results show a putative molecular basis for the hypermyelinating phenotype observed in patients with this particular mutation and provide overall information on the effects of disease-linked mutations in a flexible, membrane-binding protein segment.

## Introduction

Fast saltatory nerve impulse conduction requires myelin, a structure composed of tightly stacked lipid bilayers that wrap around selected axonal segments in the central and peripheral nervous systems (CNS and PNS, respectively). The insulative nature of myelin enables efficient nerve impulse propagation, and the destruction of myelin, demyelination, underlies a range of chronic diseases. In the PNS, peripheral neuropathies affect Schwann cell compact myelin. These include Charcot-Marie-Tooth disease (CMT) and its more severe, rapidly progressive form known as Dejerine-Sottas syndrome (DSS), which cause incurable chronic disability (Hartline 2008; Stassart *et al.* 2018). CMT and DSS manifest through both dominant and recessive inheritance, and they harbour a strong genetic component, typically caused by mutations in proteins relevant for the formation and stability of PNS myelin, while axonal forms also exist.

Myelin protein zero (P0) is a type I transmembrane protein consisting of an extracellular immunoglobulin (Ig)-like domain (Shapiro *et al.* 1996), a single transmembrane helix, and a 69-residue C-terminal cytoplasmic tail (P0ct). P0ct is likely to be involved in the regulation of myelin membrane behaviour, supporting the arrangement of the P0 Ig-like domains in the extracellular space upon the formation of the myelin intraperiod line (Luo *et al.* 2007; Raasakka *et al.* 2019b; Wong and Filbin 1994). P0ct contains a neuritogenic segment, which can be used to induce experimental autoimmune neuritis (EAN) in animal models (de Sèze *et al.* 2016). *In vitro*, P0ct is disordered in aqueous solution, gaining secondary structure upon binding to negatively charged phospholipids (Luo *et al.* 2007; Raasakka *et al.* 2019b). In its lipid-bound state, P0ct affects the phase behaviour of lipids and promotes the fusion of lipid vesicles. High-degree molecular order, most likely from stacked lipid bilayers, can be detected *via* X-ray diffraction of P0ct-bound membranes (Raasakka *et al.* 2019b). This suggests that P0ct harbours a structural role in mature myelin.

Dozens of mutations have been identified in P0, most of which affect the Ig-like domain. These mutations affect myelin morphology and integrity, leading to the development of peripheral neuropathies (Mandich *et al.* 2009; Shy *et al.* 2004). Six known missense mutations are located within P0ct, of which four cause dominant demyelinating CMT type 1B (CMT1B). These include T216ER (Su *et al.* 1993), D224Y (also referred to as D195Y and D234Y) (Fabrizi *et al.* 2006; Miltenberger-Miltenyi *et al.* 2009; Schneider-Gold *et al.* 2010), R227S (Shy *et al.* 2004), and the deletion of Lys236 (K236del) (Street *et al.* 2002). In addition, K236E has been linked to dominant axonal CMT type 2I (CMT2I) (Choi *et al.* 2004), and A221T, which was discovered as a co-mutation together with the deletion of Val42 in the Ig-like domain, was identified in a patient with DSS (Planté-Bordeneuve *et al.* 2001). How these mutations relate to CMT/DSS etiology is not known, although P0 mutations have been linked to the unfolded protein response (UPR) (Bai *et al.* 2013; Bai *et al.* 2018; Wrabetz *et al.* 2006), indicating issues in either translation or folding that induce stress within the endoplasmic reticulum (ER).

Considering the small size of P0ct and the nature of the disease mutations in it, many of which change its electrostatic charge, impairment in the function of P0ct as a membrane binding/stabilizing segment is a possible functional mechanism. We used methodologies established earlier for myelin basic protein (MBP) (Raasakka *et al.* 2017) and wild-type P0ct (wt-P0ct) (Raasakka *et al.* 2019a; Raasakka *et al.* 2019b) to characterize structure-function relationships of the CMT- and DSS-related P0ct variants. Our results suggest that D224Y is a hypermyelinating gain-of-function mutation, which is in line with the clinically relevant phenotype of abnormally thickened myelin sheaths (Fabrizi *et al.* 2006).

## Results

We have earlier studied the binding of MBP and P0ct to model lipid membranes (Raasakka *et al.* 2017; Raasakka *et al.* 2019a; Raasakka *et al.* 2019b), using a biophysical workflow that allows the determination of binding affinity, gain in folding, alteration of lipid phase behaviour, quantification and visualization of vesicle aggregation and fusion, and supported lipid bilayer (SLB) stacking. In the current study, we examined whether and how CMT and DSS mutations within P0ct influence its structure and function. For this purpose, we expressed and purified the wild-type protein and six mutant variants, each harbouring one of the following amino acid changes: T216ER, A221T, D224Y, R227S, K236E, and K236del.

### Characterization of P0ct CMT mutants

wt-P0ct and the six CMT variants were purified to homogeneity. Most mutants were straightforward to purify, showing identical behaviour to wt-P0ct in size-exclusion chromatography (SEC) (Fig. 1b). D224Y, on the other hand, had to be gel filtrated at a higher pH and salt concentration than the others, and while yields were generally lower, minor amounts of degradation were present and the migration in SEC was altered, albeit not in denaturing gel electrophoresis (SDS-PAGE) (Fig. 1b. Supplementary Fig. S1). In dynamic light scattering (DLS), all variants displayed a similar hydrodynamic radius (*R*_h_) and an absence of aggregation (Fig. 1c, Supplementary Table 1). All of the variants showed high apparent molecular weight in SDS-PAGE, which reflects the intrinsically disordered nature of P0ct (Raasakka *et al.* 2019b). The molecular weight and the presence of the mutations were confirmed using mass spectrometry (Table 1). The total yields of the purified mutant proteins were different from wt-P0ct (Supplementary Fig. S1, Table 1), most mutants giving larger yields, with the exception of D224Y. It should be noted that all mutants were expressed as maltose-binding protein fusions. Thus, mutations, which represent small changes in the overall sequence and size of the fusion protein, can affect the expression and purification behaviour.

**Fig. 1.**
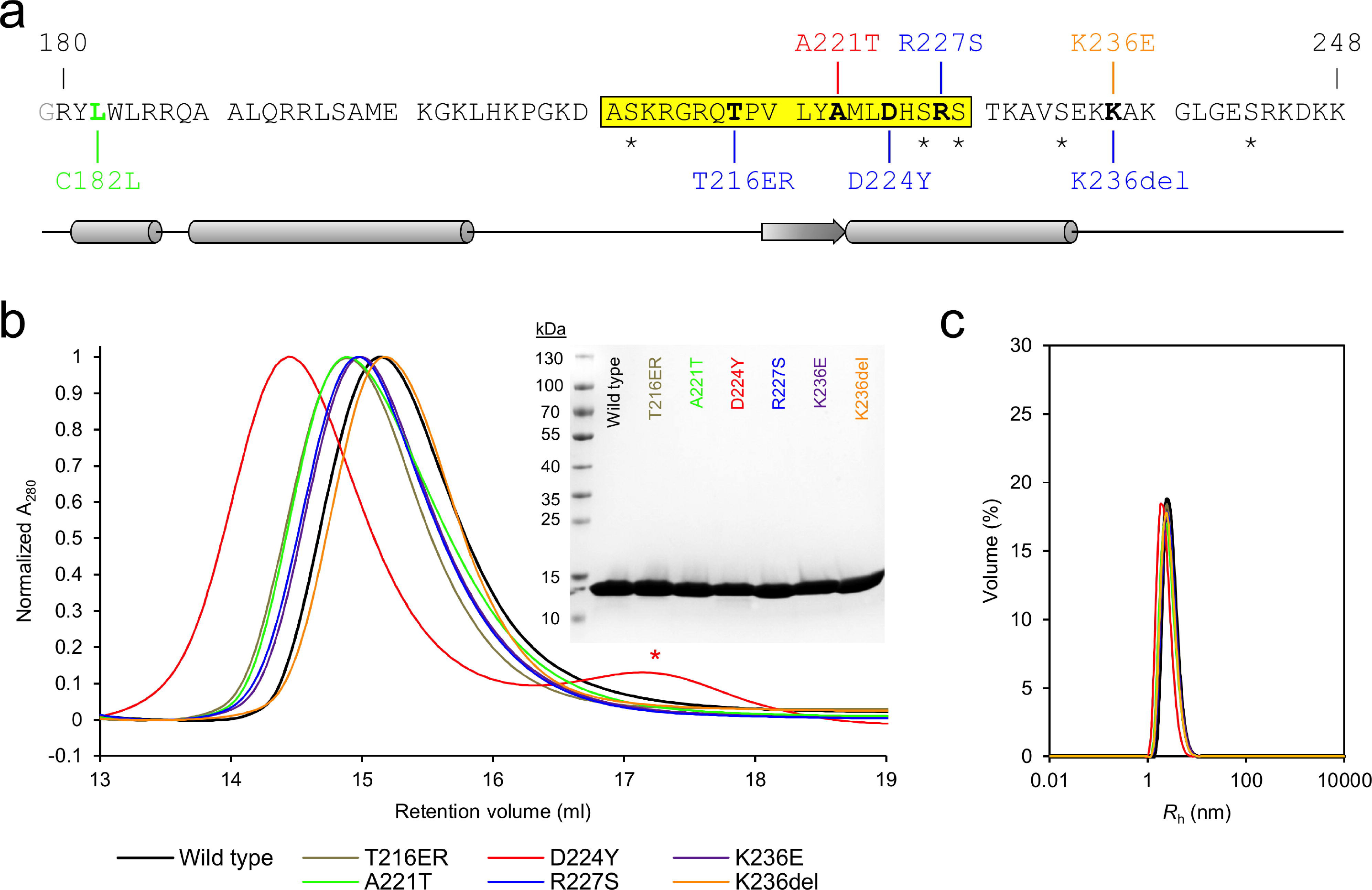
Overview of P0ct mutations. (a) The primary sequence of the wt-P0ct construct, corresponding to amino acids 180 – 248 of the human P0 precursor, with an extra N-terminal Gly residue (gray) left behind from affinity tag cleavage. The Cys182 palmitoylation site was mutated into a Leu (green) in all constructs. Putative serine phosphorylation sites are indicated with asterisks. Residues affected by disease mutations are in bold. CMT1B, CMT2I, and DSS point mutations are shown in blue, red, and orange, respectively. The sequence highlighted in yellow corresponds to the neuritogenic segment used in EAN models (de Sèze *et al.* 2016). Secondary structure prediction is shown below. (b) SEC traces of wt-P0ct and mutants as determined using a Superdex 75 10/300GL column. Note the slightly earlier retention volume of D224Y, for which the chromatography had to be performed with a different running buffer than for the other variants. The degradation products (red asterisk) present with D224Y could be completely removed using SEC. The final purity of each P0ct variant (4 µg per lane) as determined using SDS-PAGE is shown as inset. (c) DLS data of P0ct variants display good monodispersity with minimal variation in *R*_h_.

**Table 1.** Recombinant protein characterization.

| Variant* | Condition | pI** | Purification |  | Molecular weight determination |  |  | Peptide fingerprinting |
| --- | --- | --- | --- | --- | --- | --- | --- | --- |
|  |  |  | Yield*** | Solubility | Measured | Theoretical** | Difference | Mutation confirmed |
| wt-P0ct | - | 11.11 | 2.1 ± 0.4 | ++ | 7989.0 | 7990.35 | -1.35 | - |
| T216ER | CMT1 | 11.08 | 4.2 ± 0.4 | +++ | 8173.0 | 8174.54 | -1.54 | yes |
| A221T | DSS | 11.11 | 5.0 ± 0.7 | +++ | 8018.0 | 8020.37 | -2.37 | yes |
| D224Y | CMT1 | 11.12 | 0.8 ± 0.3 | + | 8037.0 | 8038.43 | -1.43 | yes |
| R227S | CMT1 | 10.89 | 6.1 ± 2.0 | +++ | 7919.0 | 7921.24 | -2.24 | yes |
| K236E | CMT2 | 10.85 | 5.1 ± 1.8 | +++ | 7989.0 | 7991.29 | -2.29 | yes |
| K236del | CMT1 | 11.09 | 5.2 ± 1.0 | +++ | 7860.0 | 7862.17 | -2.17 | yes |

\* All proteins, including wt-P0ct, contain the C182L mutation.
\*\* Values determined from protein sequences using ProtParam
\*\*\* Expressed as mg protein obtained on average per liter culture. See Supplementary Fig. 1 for graphical representation.

Small-angle X-ray scattering (SAXS) verified that for most variants, both the size and behaviour in solution were nearly identical, with radius of gyration (*R*_g_) and maximum dimension (*D*_max_) at 2.4 - 2.7 nm and 9.0 –10.7 nm, respectively, and molecular masses matching monomeric protein based on *I*_0_ values (Fig. 2, Supplementary Table 2). D224Y presented a marginally larger *D*_max_ (11.6 nm) compared to the other variants, but all variants were flexible and extended in solution, as evident from the Kratky plot (Fig. 2d).

**Fig. 2.**
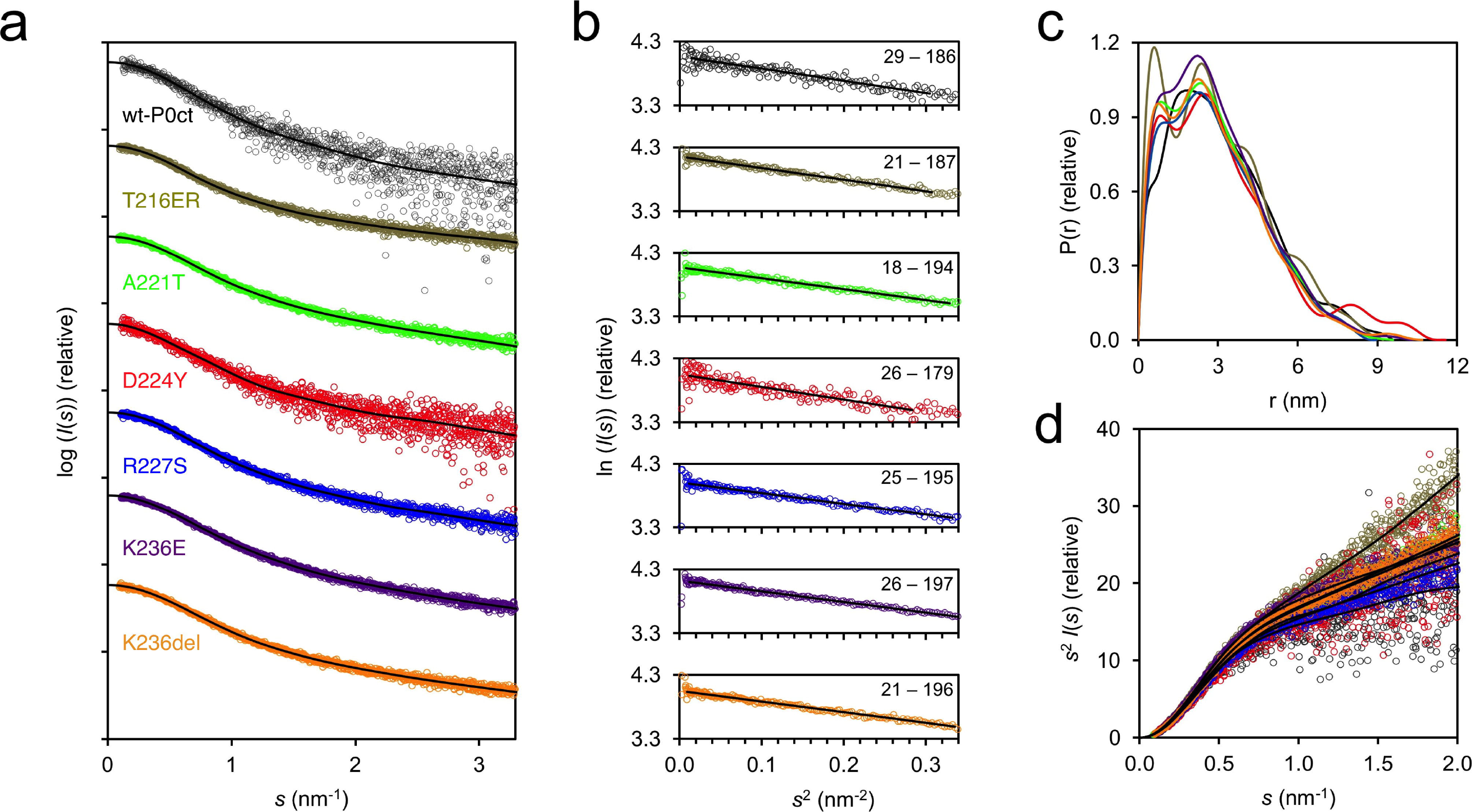
SAXS analysis of P0ct in solution. (a) SAXS data for P0ct variants. The scattering curves have been offset for clarity. (b) Guinier fits based on SAXS data. Data range is shown within each graph. (c) Distance distributions. (d) Kratky plots. P0ct variant data point coloring is consistent throughout the figure. GNOM fits to the data are shown as black lines in panels (a) and (c).

### The folding and lipid binding properties of P0ct CMT mutants

To compare the conformation of the P0ct variants, we carried out a series of synchrotron radiation circular dichroism (SRCD) spectroscopic experiments in the absence and presence of different lipid compositions, detergents, and 2,2,2-trifluoroethanol (TFE), as previously described for wt-P0ct (Raasakka *et al.* 2019b). P0ct is disordered in solution and gains a significant amount of secondary structure upon binding to small unilamellar vesicles (SUV) with a net negative surface charge (Luo *et al.* 2007; Raasakka *et al.* 2019b). In water, all mutants were disordered as expected, with D224Y having less secondary structure than the others (Fig. 3). This is in agreement with the longer *D*_max_ determined using SAXS. All mutants closely resembled wt-P0ct in TFE and the detergents sodium dodecyl sulphate (SDS), *n*-dodecylphosphocholine (DPC), lauryldimethylamine *N*-oxide (LDAO), and *n*-octyl glucoside (OG), while K236del was more α-helical than the other variants in the presence of SDS (Fig. 3, Supplementary Fig. S2).

**Fig. 3.**
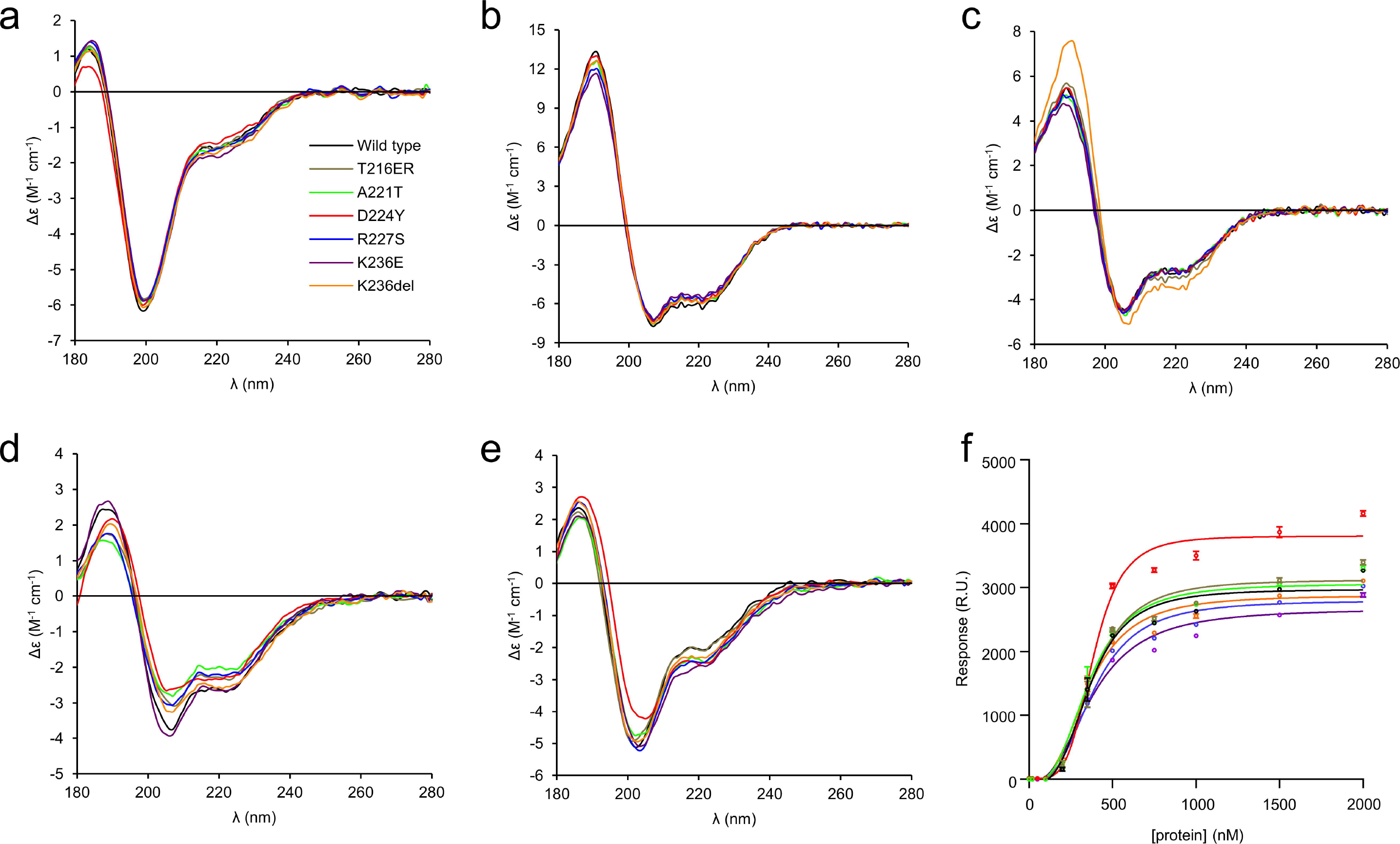
Folding and lipid binding analysis of P0ct variants. The folding of wt-P0ct and mutants was studied using SRCD spectropolarimetry in (a) water, (b) 30% TFE, (c) 0.5% SDS, (d) DMPC:DMPG (1:1), and (e) DMPC:DMPG (4:1) at 1:200 P/L ratio in each lipid condition. Additional spectra are presented in Supplementary Fig. S2. (f) SPR measurements were used to determine the affinity of each P0ct variant to immobilized DMPC:DMPG (1:1) vesicles. The colour coding legend in panel (a) for each mutant trace also corresponds to all other traces in subsequent panels.

Addition of DMPC retained the proteins in a disordered state, with D224Y deviating slightly (Supplementary Fig. S2). In the presence of net negatively charged SUVs composed of DMPC:DMPG ratios of 1:1, 4:1, and 9:1, the variants presented some folding differences (Fig. 3, Supplementary Fig. S2). Overall, most folding was observed in 1:1 DMPC:DMPG, and the degree of folding decreased with decreasing fraction of DMPG, *i.e.* negative charge. In DMPC:DMPG (1:1), a small shift to the right of the maximum at 188 nm was evident for D224Y and K236del, indicating slightly increased folding, although the two minima at 208 and 222 nm, typical for helical content, remained the same for all variants (Fig. 3d). In DMPC:DMPG (4:1), this effect was only observed for D224Y (Fig. 3e). In DMPC:DMPG (9:1), the differences in signal magnitude were large, reflecting different levels of turbidity (Supplementary Fig. S2). It can be assumed that the variants showing high turbidity under this condition are membrane-bound, while the ones giving strong CD signal of an unfolded protein do not bind to 9:1 DMPC:DMPG.

The affinity of P0ct variants towards immobilized DMPC:DMPG (1:1) SUVs was investigated using surface plasmon resonance (SPR). All variants bound to lipids with similar kinetic parameters (Fig. 3f, Table 2), including the *A*_1_ value, which corresponds to the apparent *K*_d_, of 0.35-0.4 µM. This value in the same range with those obtained earlier for wt-P0ct, MBP, and P2 (Raasakka *et al.* 2017; Raasakka *et al.* 2019b; Ruskamo *et al.* 2014; Wang *et al.* 2011). While the differences in *K*_d_ were minor, the behaviour of D224Y was unique: the observed maximal response level was higher compared to the other variants. This suggests that the D224Y variant can either accumulate onto immobilized vesicles in higher amounts, or it induces a change on the surface that affects the measurement, such as the fusion, swelling, or aggregation of lipid vesicles.

**Table 2.** SPR fitting parameters.

| Variant | R<sub>hi</sub> | R<sub>lo</sub> | A<sub>1</sub> | A<sub>2</sub> | R<sup>2</sup> |
| --- | --- | --- | --- | --- | --- |
| wt-P0ct | 2975 ± 79 | -69.10 ± 61.54 | 363.2 ± 15.1 | 3.237 ± 0.411 | 0.9858 |
| T216ER | 3123 ± 86 | -44.63 ± 64.19 | 375.1 ± 15.6 | 3.173 ± 0.409 | 0.9854 |
| A221T | 3061 ± 82 | -44.81 ± 62.30 | 357.0 ± 15.5 | 2.973 ± 0.363 | 0.9863 |
| D224Y | 3811 ± 81 | 11.30 ± 66.26 | 385.2 ± 11.3 | 4.416 ± 0.540 | 0.9886 |
| R227S | 2798 ± 78 | -39.52 ± 55.20 | 384.9 ± 16.0 | 2.936 ± 0.361 | 0.9864 |
| K236E | 2671 ± 92 | -49.48 ± 60.00 | 380.8 ± 20.0 | 2.526 ± 0.340 | 0.9831 |
| K236del | 2880 ± 79 | -33.85 ± 58.90 | 356.1 ± 16.0 | 2.852 ± 0.347 | 0.9862 |

### Effect of CMT mutations on lipid membrane properties

To determine the effect of the mutations on lipid structure, experiments probing changes in the thermodynamic and structural properties of lipid membranes were carried out. As shown before (Raasakka *et al.* 2019b), the presence of P0ct changes the melting behaviour of dimyristoyl lipid tails, inducing a population that melts 0.9 °C below the major phase transition temperature of 23.8 °C. The presence of the mutations altered this effect mildly (Fig. 4a), with T216ER and R227S behaving similarly to wt-P0ct. The Lys236 mutations deviated from wt-P0ct, with a decreased temperature for the emerged population; K236E and K236del showed lipid phase transition temperatures of 22.7 and 22.8 °C, respectively. A221T presented slightly higher temperature for phase transition compared to wt-P0ct, with the major peak at 23.0 °C. Based on the shape of the calorimetric landscape, D224Y was clearly different from the rest, as the new population did not appear as a single, symmetric peak, but was rather formed of several overlapping peaks.

**Fig. 4.**
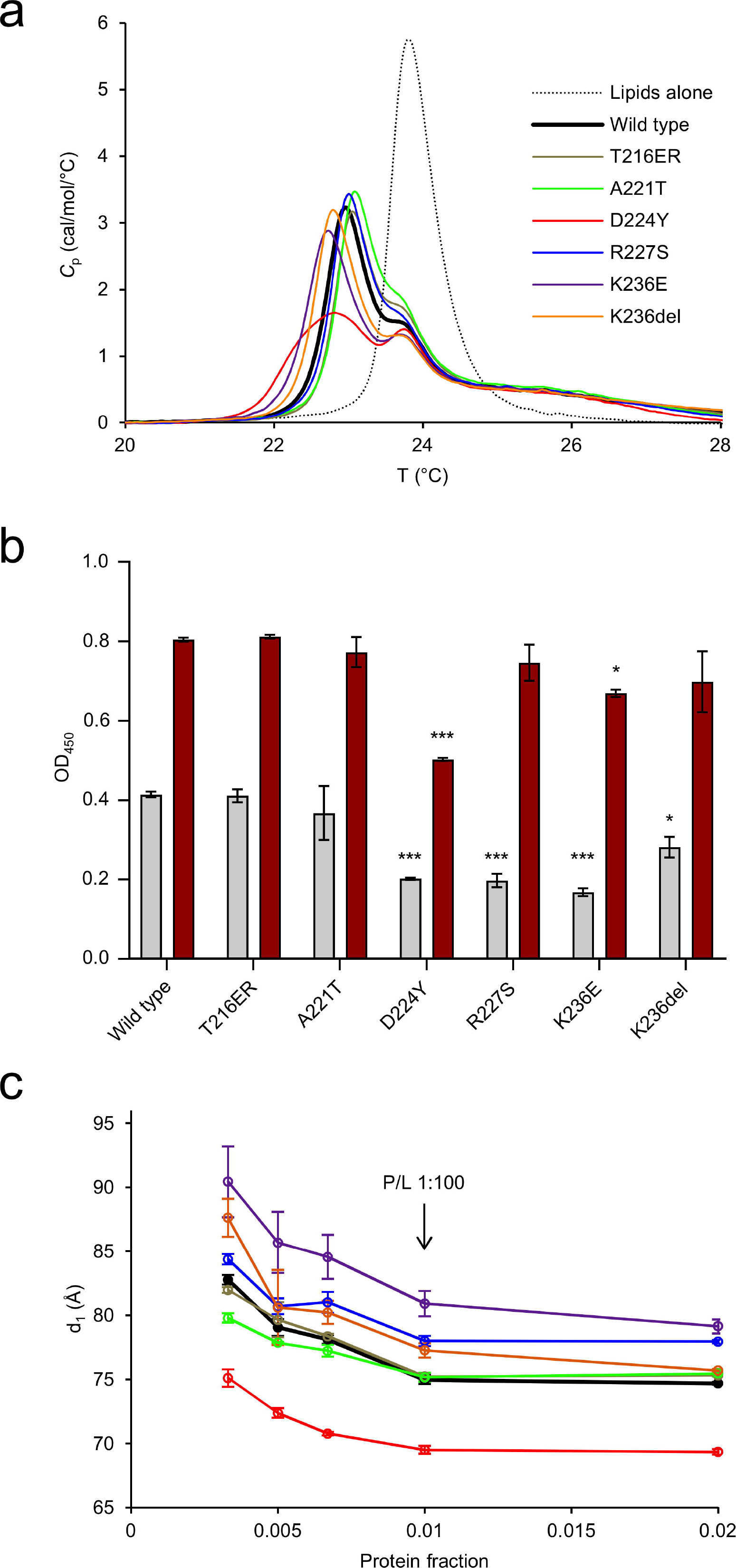
Analysis of protein-induced lipid structure behavior. (a) DSC analysis of lipid tail transition behaviour. (b) Turbidimetric analysis of 0.5 mM DMPC:DMPG (1:1) at 5 µM (gray) and 10 µM protein concentration (dark red). These proteins concentrations translate to 1:100 and 1:50 P/L ratios, respectively. Error bars represent standard deviation. Statistical analysis was performed using one-way analysis of variance (ANOVA) followed by Dunnett’s multiple comparisons test to wt-P0ct turbidity within the same protein concentration series (* : P < 0.05; *** : P < 0.001). (c) SAXD analysis reveals that D224Y displays a significantly tighter mean repeat distances compared to wt-P0ct, whereas K236E is most loose. The traces have identical colouring to (a).

Similarly to MBP and P2 (Raasakka *et al.* 2017; Ruskamo *et al.* 2014), P0ct is capable of inducing concentration-dependent solution turbidification, when mixed with lipid vesicles of net negative charge (Raasakka *et al.* 2019b). The turbidity can arise from vesicle fusion and/or aggregation, and different processes may be dominant in different samples with respect to the measured signal. To determine the effect of P0ct CMT mutations on this function, turbidity experiments were carried out with the different variants. T216ER and A221T produced turbidity levels similar to wt-P0ct (Fig. 4b, Supplementary Fig. S3a). At 1:100 P/L ratio, D224Y, R227S, K236E, and K236del all had decreased turbidity. At a P/L ratio of 1:50, however, only D224Y had a significant inhibitory effect on turbidity. This result highlights that the D224Y mutant protein may function differently from the other variants, when it binds to and aggregates vesicles.

To shed further light on the protein-induced changes in membrane structure, small-angle X-ray diffraction (SAXD) experiments were performed on P0ct-membrane mixtures. In our earlier study, wt-P0ct mixed with lipids produced two strong Bragg peaks, and the corresponding repeat distance evolved as a function of the P/L ratio (Raasakka *et al.* 2019b). Here, we observed that in all cases, the repeat distance increased when protein concentration in the sample decreased (Fig. 4c, Supplementary Fig. S3b). Each variant presented a minimum repeat distance, which was reached at and above a P/L ratio of 1:100. The repeat distance for wt-P0ct was ~7.5 nm, while D224Y produced a spacing of <7.0 nm. R224S, K236E, and K236del had looser packing than wt-P0ct. K236E had a minimum repeat distance of ~8.0 nm at the highest protein concentration.

To understand the effect of the mutations on the function of P0ct, and the origin of the high molecular order reflected by X-ray diffraction, electron microscopy imaging was performed. Most mutants functioned in a manner similar to wt-P0ct, producing large vesicular structures with a spread-out morphology (Fig. 5), with occasional regions indicative of bilayer stacking. D224Y showed a clear difference to wt-P0ct, producing strongly stacked myelin-like membranes in a manner resembling MBP (Raasakka *et al.* 2017). This gain of function was reproducible over a wide range of P/L ratios (Supplementary Fig. S4) and a unique feature among the six mutant P0ct variants. The results confirm that the Bragg peaks seen in SAXD, indeed, originate from repeat distances in membrane multilayers, identically to two other PNS myelin peripheral membrane proteins, MBP and P2 (Raasakka *et al.* 2017; Ruskamo *et al.* 2014; Sedzik *et al.* 1985). The observed bilayer spacing for the D224Y mutant in EM was narrow and in general better defined than seen for MBP (Raasakka *et al.* 2017), suggesting that P0ct forms a tight structure within and/or between the membranes. Based on SAXD, the intermembrane spacing is ~3 nm, a value in close relation to the dimensions of the major dense line (MDL) in myelin.

**Fig. 5.**
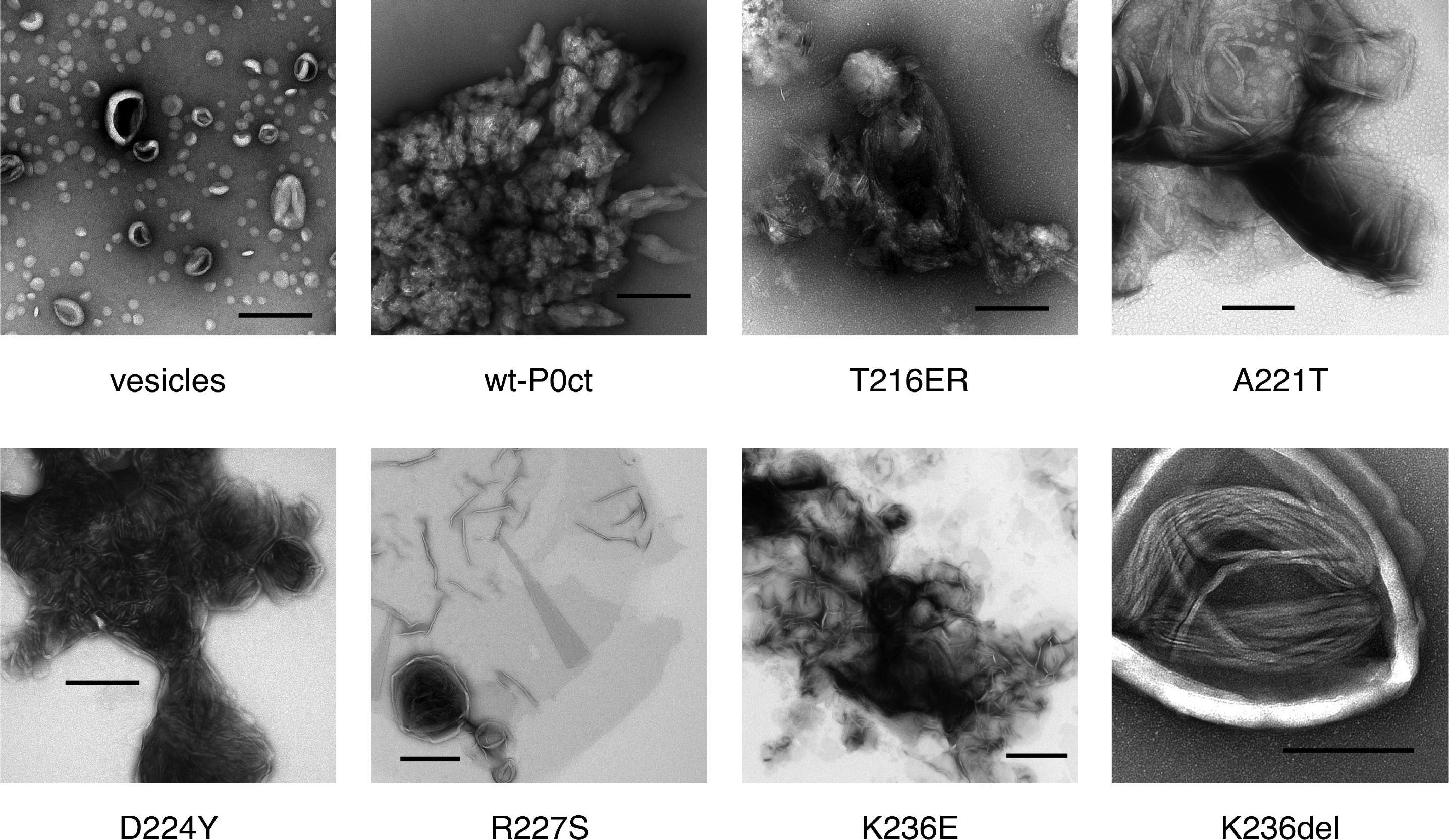
EM analysis of P0ct mutants. Negatively stained samples of DMPC:DMPG (1:1) vesicles (a) alone, and (b) wt-P0ct, (c) T216ER, (d) A221T, (e) D224Y, (f) R227S, (g) K236E, and (h) K236del were imaged at 1:200 P/L ratio. D224Y forms multilayered lipid structures that are absent for wt-P0ct.

To gain an insight into the kinetic aspects of P0ct-induced lipid fusion/aggregation, stopped-flow kinetics experiments were performed using SRCD (Fig. 6, Table 3) (Raasakka *et al.* 2019a). All variants followed a similar kinetic pattern as wt-P0ct and could be best fitted to a two-phase exponential decay with two rate constants (*k*_1_, fast and *k*_2_, slow). Rather minor differences were present: *k*_2_ values were very similar in all cases, and while D224Y presented 10% higher *k*_1_ and *k*_1_/*k*_2_ compared to wt-P0ct and most other variants, both K236E and K236del displayed *k*_1_ and *k*_1_/*k*_2_ 20% lower than for wt-P0ct, indicating slower kinetics (Fig. 6b). While all variants produced a similar end-level CD value around −100 mdeg, the starting level of K236del was higher than for any other variant, and remained so until ~0.3 s, before settling on a similar level to other variants. It is currently unclear whether this is due to an increased level of protein folding or less scattered light from fused or aggregated vesicles.

**Fig. 6.**
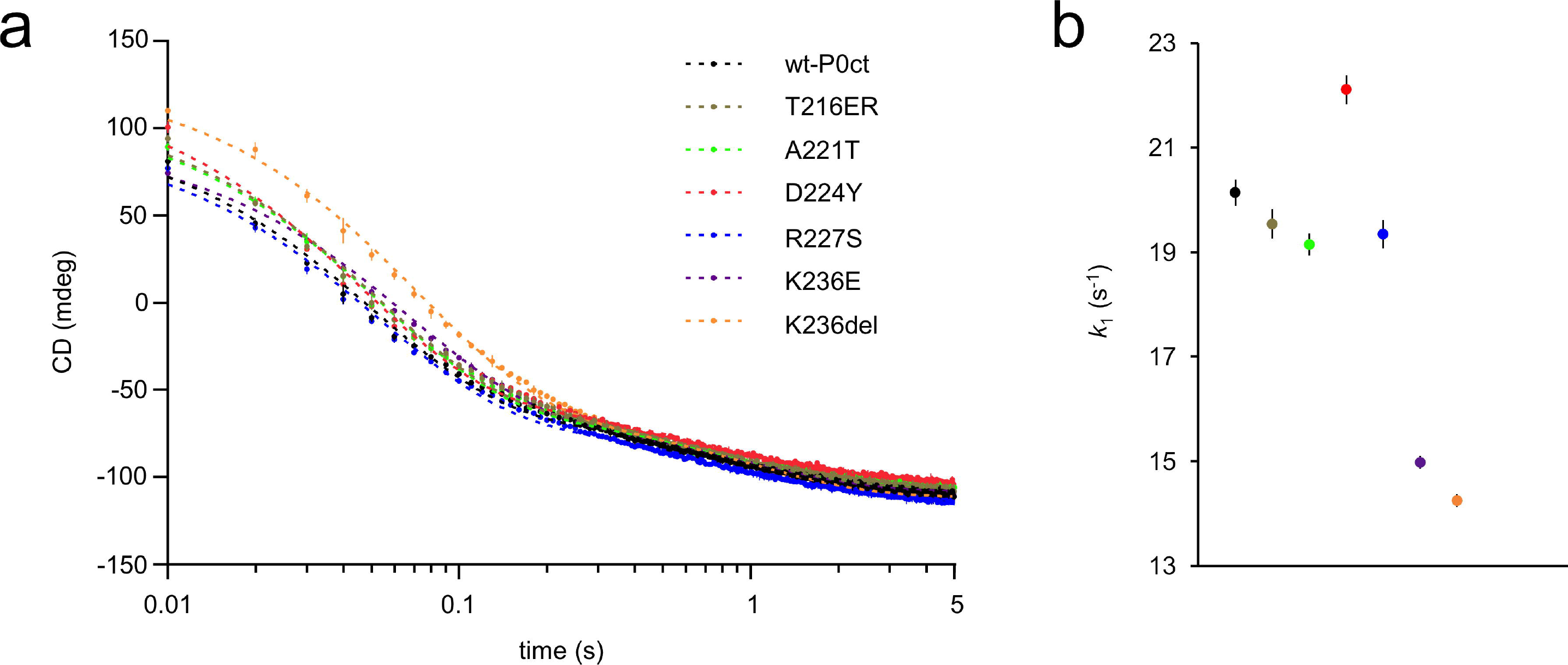
SRCD stopped-flow kinetics of protein-induced initial lipid turbidification. (a) The SRCD signal evolution was monitored using rapid kinetics at 195 nm for 5 sec. wt-P0ct and mutants were mixed with DMPC:DMPG (1:1) lipids at 1:200 P/L ratio in the presence of 150 mM NaF. Fits (dashed lines) are plotted over the measurement points. Error bars represent standard deviation. See Table 4 for fitting results. (b) Graphical comparison of the obtained *k*_1_ values.

**Table 3.** Kinetic constants for protein-induced vesicle turbidity. The kinetic constants were obtained by fitting the data to a two-phase exponential decay function. All errors represent standard deviation.

| Variant | k<sub>1</sub> (s<sup>-1</sup>) | k<sub>2</sub> (s<sup>-1</sup>) | k<sub>1</sub>/k<sub>2</sub> | R<sup>2</sup> |
| --- | --- | --- | --- | --- |
| wt-P0ct | 20.14 ± 0.25 | 1.12 ± 0.01 | 17.96 ± 0.22 | 0.9934 |
| T216ER | 19.54 ± 0.28 | 1.18 ± 0.02 | 16.63 ± 0.25 | 0.9908 |
| A221T | 19.15 ± 0.21 | 1.14 ± 0.01 | 16.76 ± 0.20 | 0.9943 |
| D224Y | 22.11 ± 0.28 | 1.19 ± 0.02 | 18.53 ± 0.25 | 0.9923 |
| R227S | 19.35 ± 0.27 | 1.08 ± 0.02 | 17.95 ± 0.26 | 0.9916 |
| K236E | 14.98 ± 0.12 | 1.02 ± 0.01 | 14.64 ± 0.13 | 0.9969 |
| K236del | 14.25 ± 0.12 | 1.05 ± 0.01 | 13.54 ± 0.13 | 0.9967 |

### The membrane insertion mode of P0ct

To understand the membrane insertion of P0ct, how it compares to MBP (Raasakka *et al.* 2017), and how it might be related to disease mutations, we performed neutron reflectometry (NR) experiments (Fig. 7, Table 4). The insertion of P0ct to a DMPC:DMPG SLB was quite different to that of MBP. P0ct inserted completely into the membrane, thickening it by 2 nm and increasing its roughness, most likely due to increased bilayer mobility, as the hydration layer below the membrane became thicker (Fig. 7b,c). P0ct was present in the acyl tail fraction of the membrane, as well as the outer headgroup fraction. The data could not be fitted with only these parameters, but a very rough, narrow layer of protein had to be considered on top of the membrane. Unfortunately, the roughness and high solvation fraction of this layer did not allow for precise thickness determination: the layer was modelled to be between 5 – 15 Å thick within the fit to the data. To investigate the effect of the D224Y mutation on P0ct membrane association, NR data were collected for SLB-bound D224Y, which appeared identical to wt-P0ct (Supplementary Fig. S5).

**Fig. 7.**
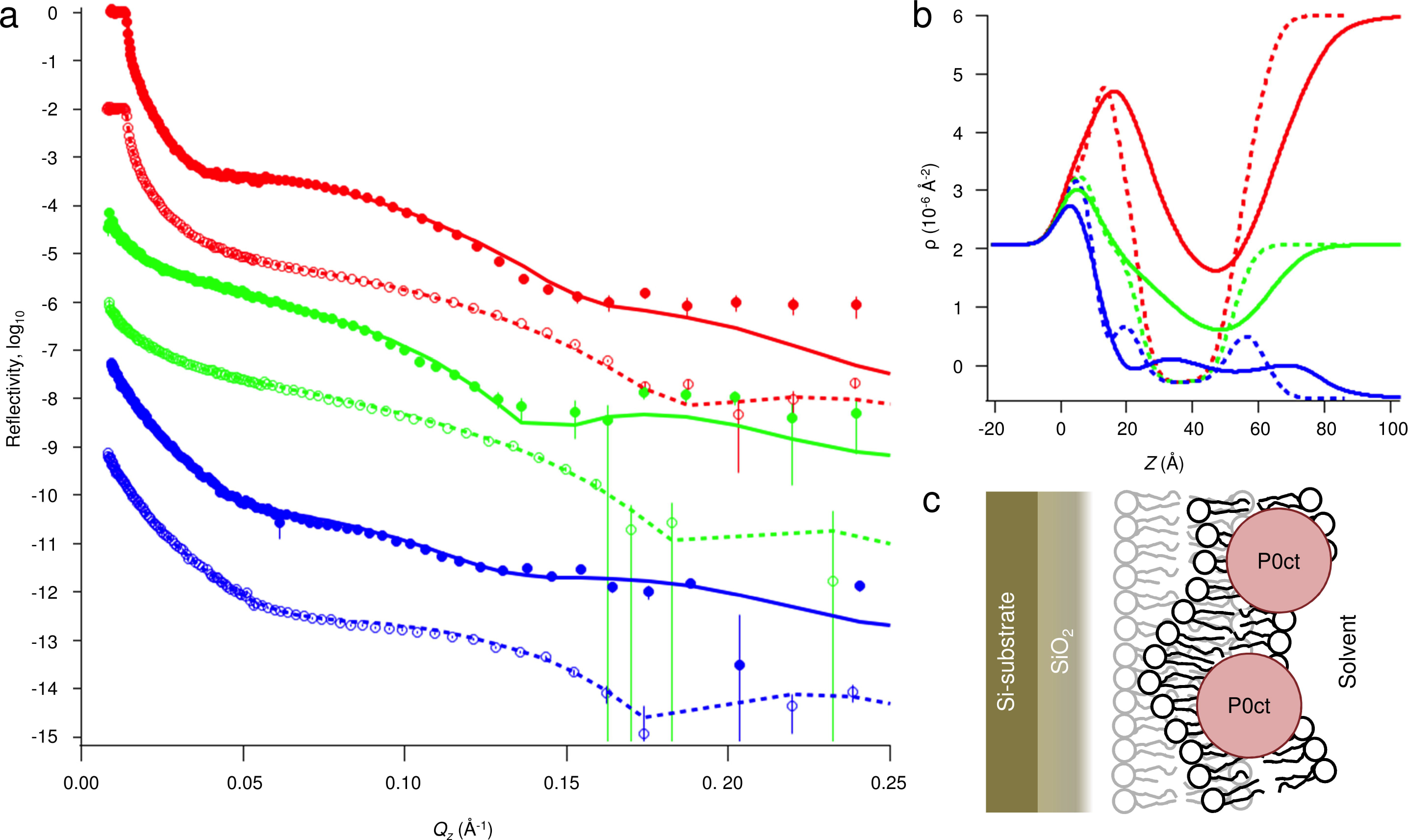
NR data and fitting. (a) NR data of a supported DMPC:DMPG (1:1) bilayer before (open markers) and after incubation with wt-P0ct (closed markers). The used solvent contrasts were 95% D_2_O (red), Si-matched water (SMW, 38% D_2_O; green) and 100% H_2_O (blue). Fits are shown as dashed and solid lines for the bilayer before and after addition of wt-P0ct, respectively. (b) Scattering length density (ρ) profiles obtained from the fitting. The error bars denote standard deviation. (c) Model for the P0ct-bound membrane. The protein-free membrane is shown in light gray on the background.

**Table 4.** NR parameters.

|  |  | **Bilayer alone** | **Bilayer with 0.5 μM wt-P0ct** |
| --- | --- | --- | --- |
| **Substrate** | Oxide thickness ($\text{\AA}$) | 10.6 | 11 |
|  | Oxide solvation (%) | 0 | 0 |
| | Oxide roughness ($\text{\AA}$) | 4 | 4 |
| | Hydration layer between oxide and bilayer ($\text{\AA}$) | 4.6 | 12 |
| | Hydration layer roughness ($\text{\AA}$) | 3 | 6 |
| **Bilayer** | Bilayer area-per-molecule ($\text{\AA}^2/\text{molecule}$) | 60 | 70 |
| | Inner headgroups thickness ($\text{\AA}$) | 8.3 | 8 |
| | Inner headgroups roughness ($\text{\AA}$) | 3.6 | 8.1 |
|  | Inner headgroups solvation (%) | 35 | 45 |
| | Acyl tails thickness ($\text{\AA}$) | 28.8 | 32 |
| | Acyl tails roughness ($\text{\AA}$) | 3.8 | 13.3 |
|  | Acyl tails hydration (%) | 0 | 17 |
| | Outer headgroups thickness ($\text{\AA}$) | 8.8 | 8 |
| | Outer headgroups roughness ($\text{\AA}$) | 4.9 | 9.5 |
|  | Outer headgroups solvation (%) | 35 | 53.5 |
| **wt-P0ct** | Protein in inner headgroups (%) |  | 0 |
|  | Protein in acyl tails (%) |  | 10 |
|  | Protein in outer headgroups (%) |  | 20 |
| | Protein layer thickness ($\text{\AA}$) | | 7 |
| | Protein layer roughness ($\text{\AA}$) | | 15 |
|  | Protein layer solvation (%) |  | 86 |

## Discussion

The formation of compact myelin and the major dense line requires an interplay of myelin molecules, many of which have similar functional properties despite lack of sequence homology. Considering the MDL of PNS compact myelin, the major protein components according to current knowledge are MBP, P2, P0ct, and cytosolic loops of PMP-22. We characterized the potential functional anomalies of P0ct CMT mutants in membrane binding using earlier established biophysical strategies (Raasakka *et al.* 2017; Raasakka *et al.* 2019a; Raasakka *et al.* 2019b).

The six mutations reported in P0ct are clustered within or near the neuritogenic segment. Most of them reside in the vicinity of putatively phosphorylated Ser residues (Fig. 1a), which have been speculated to be affected by P0ct mutations (Su *et al.* 1993; Xu *et al.* 2001). Many P0 mutations have been suggested to lead to UPR activation (Bai *et al.* 2013; Bai *et al.* 2018; Wrabetz *et al.* 2006), indicating problems with translation rate, folding, and/or membrane insertion. Given the fact that P0ct is known to interact with lipid membrane surfaces (Luo *et al.* 2007; Raasakka *et al.* 2019b; Raasakka *et al.* 2019a), these mutations could also have direct effects on the formation of mature compact myelin at the molecular level.

### Mechanism of P0ct binding to membranes

In order to fully understand the effects of P0ct mutations on its structure and function, detailed knowledge about P0ct binding to lipid membranes, and the effects thereof on multilayered membrane stacks, are required. NR allowed us to gain a picture of P0ct in a lipid bilayer. P0ct inserts deep into a membrane, with only a small fraction remaining solvent-exposed on the membrane surface. This is a clear difference to MBP, which forms a brush-like protein phase on top of the membrane surface, while being partially embedded into the bilayer (Raasakka *et al.* 2017). After undergoing charge neutralization and folding, P0ct seems to collapse into a tight conformation and remain stable. The compact, deep conformation of P0ct suggests that instead of directly embedding into two bilayers, which is the working model for *e.g.* MBP-induced stacking (Raasakka *et al.* 2017; Vassall *et al.* 2015), P0ct may change the surface properties of the membrane in a way that supports apposing bilayer surface adhesion. It could also regulate membrane curvature and the twining of lipid bilayers around the axon.

At the level of full-length P0, P0ct is a direct extension of the transmembrane segment, and hence, anchored permanently to a membrane surface at its beginning. Membrane stacking could involve the insertion of P0ct across the MDL into an apposing membrane leaflet, which is only 3 nm away. Considering this scenario, P0 is basally expressed in Schwann cells even before myelination occurs (Lee *et al.* 1997). Moreover, P0 is translated and inserted into the ER membrane and trafficked through the trans-Golgi network to the plasma membrane after the Ig-like domain has been post-translationally modified (Eichberg 2002; Lemke and Axel 1985). If P0ct were to enter an apposing membrane during the formation of compact myelin, it would have to remain in a disordered state until another membrane is present. On the other hand, if P0ct is embedded in the membrane after translation, it might afterwards be able to dissociate and enter the apposing leaflet within compact myelin. Considering the attractive phospholipid bilayer around the transmembrane helix, and the fact that P0ct binds negatively charged lipids essentially irreversibly *in vitro* ^5^, both mechanisms described above are unlikely to exist. Thus, the role of P0ct in membrane adhesion is likely to be based on altered lipid membrane properties, as opposed to MBP and P2, which directly interact with two membrane surfaces. While P2 and MBP were observed to synergistically stack lipid bilayers *in vitro* (Suresh *et al.* 2010), mice lacking both proteins formed apparently normal and functional myelin (Zenker *et al.* 2014). Hence, multiple factors must participate in the correct formation of compact myelin; these include both lipid components of the myelin membrane, different myelin proteins, as well as signalling molecules and inorganic ions. Hence, further experiments in more complex sample environments are required to decipher the details of the molecular interplay between these factors in PNS myelin MDL formation.

### P0ct mutations and membrane interactions

Compared to wt-P0ct, we observed only subtle differences for two mutants: T216ER and A221T. While T216ER behaved very similarly to wt-P0ct, its role in CMT etiology could be of another origin than related to protein-membrane binding. A221T, on the other hand, resides in the YAML-motif, which directs the trafficking of P0 (Kidd *et al.* 2006) and might compromise the function of P0 even without inducing changes in membrane binding, especially when combined with a second mutation in the extracellular domain, such as the deletion of Val42 (Planté-Bordeneuve *et al.* 2001).

Functionally the most interesting mutant studied here is D224Y, which now has been described in at least 3 studies (Fabrizi *et al.* 2006; Miltenberger-Miltenyi *et al.* 2009; Schneider-Gold *et al.* 2010). It is a gain-of-function mutant, inducing ordered lipid bilayer stacks *in vitro*, which are more tightly packed than those formed by wt-P0ct or the other variants. The results correlate well the hypermyelinating disease phenotype (Fabrizi *et al.* 2006). Neutron reflectometry produced a nearly identical result for D224Y compared to wt-P0ct, which together with the SRCD experiments indicates that the conformation of wt-P0ct and D224Y is similar in the membrane. The change of an acidic to an aromatic residue near the lipid bilayer surface most likely enables a specific interaction between surfaces that results in the observed gain of function.

P0 is the most abundant protein in PNS myelin (Greenfield *et al.* 1973; Patzig *et al.* 2011), contributing primarily to the formation of the intraperiod line (Filbin *et al.* 1990), and molecular mechanisms of D224Y-induced tight stacking could be two-fold. Firstly, with its short repeat distance – 1-2 nm smaller compared to MBP and P2 based on SAXD (Raasakka *et al.* 2019b; Ruskamo *et al.* 2014; Sedzik *et al.* 1985) – and active membrane binding, as evident from SPR, the mutant might cause size exclusion of P2 and other factors out of the cytoplasmic stack, leading to defective compact myelin maintenance. In PNS compact myelin, P2 is even more abundant in the cytoplasmic compartment than MBP, can form membrane stacks, and harbours a maintenance role in myelin homeostasis as a lipid carrier (Ruskamo *et al.* 2014; Zenker *et al.* 2014). Secondly, the tendency of D224Y to form such ordered, tight systems might affect the Ig-like domains on the extracellular side. In the hypermyelinating phenotype of D224Y patients, membrane stacking seems condensed and regular, without abnormally loosened myelin (Fabrizi *et al.* 2006). SPR indicates that more D224Y can accumulate on membranes, and full-length P0 D224Y could accumulate and tighten up within the membrane, causing also the intraperiod line to become more crowded and/or structured. The original discovery of the D224Y mutation (Fabrizi *et al.* 2006) suggested that it has a gene dosage effect, since heterozygous carriers presented little to no symptoms. Hence, the presence of wild-type P0 can rescue the effects of the mutation. Correct gene dosage of P0 is important for normal myelination in animal models as well as CMT patients (Fabrizi *et al.* 2006; Maeda *et al.* 2012; Martini *et al.* 1995; Quattrini *et al.* 1999; Speevak and Farrell 2013; Wrabetz *et al.* 2000). The molecular details of the involved mechanisms are currently lacking. Further studies on the D224Y mutation *in vitro* and *in vivo* could help in understanding molecular aspects of both normal and abnormal myelination.

Lys236 appears to be a functionally important amino acid in P0ct. In its membrane-bound state, P0ct is likely to have Lys236 close to the lipid headgroups, and altering the charge in this environment might influence folding and the global positioning of P0ct on the membrane. Indeed, a gradual effect in membrane packing was observed in SAXD; the repeat structure loosens, as residue 236 neutralizes (K236del) and turns to negative (K236E). Turbidimetry also indicated a clear effect of charge reversal at residue 236. The Lys236 mutants folded to a similar degree as wt-P0ct, which suggests that the role of Lys236 is in packing, rather than folding. This is supported by the slower kinetic parameters for Lys236 mutants in stopped-flow measurements.

Similarly to Lys236, Arg227 could harbour a role in membrane packing. In our experiments, R227S is one of the mutants that appeared to induce weaker adhesion than the wild-type protein. The mutation results in a loosened repeat structure without a major impact on protein folding. Arg227 might be involved in electrostatic anchoring of the protein to the lipid headgroups – the R227S mutation likely has low impact on ER stress and UPR, as mutated P0 correctly localizes to the plasma membrane (Lee *et al.* 2010).

### Concluding remarks

To a large extent, the P0ct CMT variants studied here perform similarly to wt-P0ct in controlled simple environments. This might differ *in vivo*, where other components are present and P0 is present in its full-length form. Our characterization is focused on protein-lipid interactions and does not take into account possible protein-protein interactions with MBP, P2, or PMP22, which might be relevant for myelination and disease phenotypes. Nevertheless, we have uncovered critical amino acids in P0 that may contribute to the formation of healthy myelin and be involved in disease mechanisms. These include Arg227, Lys236, and Asp224. Our results shed light on the molecular fundamentals of myelination in the PNS, but more comprehensive studies in biological model systems, as well as on molecular structure and dynamics of native-like myelin, are needed for deciphering the mechanisms of the P0ct mutations causing human neuropathy.

## Experimental procedures

### Bioinformatics, mutagenesis, protein expression & purification

Secondary structure prediction for P0ct was performed using JPred (Drozdetskiy *et al.* 2015). Mutations were generated in the P0ct pHMGWA expression vector (Busso *et al.* 2005; Raasakka *et al.* 2019b) by PCR using Phusion High-Fidelity DNA polymerase (Thermo Fisher Scientific) with 5′-phosphorylated primers that introduced the desired point mutations. The samples were treated with *Dpn*I (New England Biolabs) to digest template DNA and linear vectors circularized using T4 DNA ligase (New England Biolabs), followed by transformation and plasmid isolation. The presence of mutations and integrity of the constructs was verified using DNA sequencing.

Protein expression and purification were carried out in *E. coli* BL21(DE3) as described for wt-P0ct (Raasakka *et al.* 2019b), with the exception of an added amylose-resin affinity step between Ni-NTA and size-exclusion chromatography. The step was introduced to remove any contaminating maltose-binding protein tags from the tobacco etch virus protease-digested recombinant proteins. Size exclusion chromatography was carried out using Superdex S75 16/60 HiLoad and Superdex 75 10/300GL columns (GE Healthcare) with 20 mM HEPES, 150 mM NaCl, pH 7.5 (HBS) as mobile phase, with the exception of D224Y, where a 20 mM Tris-HCl, 300 mM NaCl, pH 8.5 (TBS) solution was used. The monodispersity and *R*_h_ of all proteins were checked from filtered 1 mg/ml samples using a Malvern Zetasizer ZS DLS instrument. The D224Y mutant was then dialyzed into HBS. Additionally, all proteins were dialyzed into water prior to SRCD experiments.

### Mass spectrometry

The molecular weight and identity of the purified proteins were verified by mass spectrometry. In short, the proteins were subjected to ultra-performance liquid chromatography (UPLC) coupled electrospray ionization (ESI) time-of-flight mass spectrometry in positive ion mode, using a Waters Acquity UPLC-coupled Synapt G2 mass analyzer with a Z-Spray ESI source. This allowed us to determine the undigested masses of each purified P0ct variant. Protein identity and the presence of the desired mutations were confirmed from peptides extracted after in-gel tryptic proteolysis, using a Bruker Ultra fleXtreme matrix-assisted laser desorption/ionization time-of-flight (MALDI-TOF) mass analyzer.

### Small-angle X-ray scattering

SAXS data were collected from protein samples at 0.3 – 12.9 mg/ml in HBS and TBS on the EMBL P12 beamline, PETRA III (Hamburg, Germany) (Blanchet *et al.* 2015). Monomeric bovine serum albumin (M_r_ = 66.7 kDa; *I*_0_ = 499.0) was used as a molecular weight standard. Data were processed and analyzed using the ATSAS package (Franke *et al.* 2017), and GNOM was used to calculate distance distribution functions (Svergun 1992). See Supplementary Table 2 for further details.

### Vesicle preparation

DMPC, DMPG, and DOPC were purchased from Larodan Fine Chemicals AB (Malmö, Sweden). DOPS and the deuterated d_54_-DMPC and d_54_-DMPG were purchased from Avanti Polar Lipids (Alabaster, Alabama, USA).

Lipid stocks were prepared by dissolving dry lipids in chloroform or chloroform:methanol (9:1 v/v) at 10-30 mM. Mixtures were prepared from stocks at the desired molar ratios, followed by solvent evaporation under a stream of nitrogen and lyophilizing overnight at −52 °C. The dried lipids were stored at −20 °C or used directly for liposome preparation.

Liposomes were prepared by mixing dried lipids with water or HBS at 10-15 mM, followed by inverting at ambient temperature for at least 3 h. Multilamellar vesicles (MLV) were prepared by freeze-thaw cycles in liquid N_2_ and a warm water bath and vortexing. The cycle was performed 7 times in total. Large unilamellar vesicles (LUV) were prepared by passing fresh MLVs through a 0.1-µm membrane 11 times at 40 °C. SUVs were prepared by ultrasonication of fresh MLVs using a probe tip sonicator (Sonics & Materials Inc. Vibra-Cell VC-130) until clarified. All lipid preparations were immediately used in experiments.

### Synchrotron radiation circular dichroism spectroscopy

SRCD spectra were collected from 0.1 – 0.5 mg/ml protein samples in water on the AU-CD beamline at ASTRID2 synchrotron (ISA, Aarhus, Denmark). Samples containing lipids were prepared right before measurement by mixing proteins (P/L ratio 1:200) with SUVs. 100-µm pathlength closed circular cells (Suprasil, Hellma Analytics) were used for the measurements. Spectra were recorded from 170 to 280 nm at 30 °C. Baselines were subtracted and CD units converted to ∆∊ (M^-1^ cm^-1^) in CDtoolX (Miles and Wallace 2018). SDS and TFE were from Sigma-Aldrich and the detergents LDAO, OG, DM, and DPC from Affymetrix.

Rapid kinetic SRCD data were collected as described (Raasakka *et al.* 2019a). In short, an SX-20 stopped-flow instrument (Applied Photophysics) mounted on the AU-rSRCD branch line of the AU-AMO beamline at ASTRID2 (ISA, Aarhus, Denmark) at was used for data collection at 10 °C. 1-to-1 mixing of a 0.1 mg/ml protein solution and a DMPC:DMPG (1:1) SUV solution (at P/L ratios 1:200) was achieved using a mixer (2 ms dead time) before injection into the measurement cell (160 µl total volume, 2-mm pathlength) per shot. The CD signal (mdeg) was monitored at a fixed wavelength of 195 nm for 5 s with a total of 5 – 10 repeat shots per sample, which were averaged into a single curve. Each sample was prepared and measured in duplicate. Water baselines were subtracted from sample data. The data were fitted to different exponential functions using GraphPad Prism 7.

### Surface plasmon resonance

SPR was performed on a Biacore T200 system (GE Healthcare). According to the manufacturer’s instructions, 100-nm LUVs of 1 mM DMPC:DMPG (1:1) were immobilized on an L1 sensor chip (GE Healthcare) in HBS, followed by the injection of protein solutions. Chip regeneration was performed using a 2:3 (v:v) mixture of 2-propanol and 50 mM NaOH. The protein concentration was 20 – 2000 nM in HBS, and a single concentration per lipid capture was studied; all samples were prepared and measured in duplicate. In each run, one sample was measured twice to rule out instrumental deviation. The binding response as a function of protein concentration was plotted and fitted to the 4-parameter model,

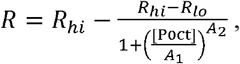

to gain information about association affinity.

### Differential scanning calorimetry

Proteins were mixed with MLVs in HBS at a protein-to-lipid ratio of 1:100 or 1:250, always containing 350 µM of DMPC:DMPG (1:1) in a final volume of 700 µl. Lipid samples without proteins were prepared as controls. The samples were degassed for 10 min under vacuum with stirring at 10 °C before measurements. DSC was performed using a MicroCal VP-DSC calorimeter with a cell volume of 527.4 µl. The reference cell was filled with HBS. Each sample was scanned from 10 to 40 °C and back to 10 °C in 1 °C/min increments. Baselines were subtracted from sample curves and zeroed between 15 & 20 °C to enable straightforward comparison between samples. All samples were prepared and measured twice, with the observed trends being reproducible.

### Vesicle turbidimetry and X-ray diffraction experiments

For turbidimetric measurements, SUVs of DMPC:DMPG (1:1) were mixed with 0.5 – 10 µM protein in duplicate and mixed thoroughly. Turbidity was recorded at 450 nm at 30 °C using a Tecan Spark 20M plate reader. Turbidity of protein-free SUVs was subtracted from the protein samples, and statistical analysis was performed using GraphPad Prism 7.

SAXD experiments were performed to investigate repetitive structures in the turbid samples. 10 and 20 µM proteins were mixed with SUVs of 1 – 3 mM DMPC:DMPG (1:1) in HBS at ambient temperature and exposed at 25 °C on the EMBL P12 BioSAXS beamline, DESY (Hamburg, Germany). A HBS buffer reference was subtracted from the data. Lipid samples without added protein did not produce Bragg peaks. The peak positions of momentum transfer, *s*, in all measured samples were used to calculate mean repeat distances, *d*, in proteolipid structures, using the equation

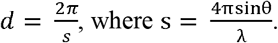

### Electron microscopy

For negatively stained EM, 740 µM DMPC:DMPG (1:1) SUVs were mixed with proteins using protein-to-lipid ratios of 1:58, 1:100, 1:200, and 1:500 and incubated at 22 °C for 1 h. EM grids were then prepared, stained and imaged as described before (Raasakka *et al.* 2017; Raasakka *et al.* 2019b; Ruskamo *et al.* 2017).

### Neutron reflectometry

Supported lipid bilayers were prepared onto flat (5 Å RMS roughness tolerance) 80 mm × 50 mm ×15 mm Si-crystal blocks (Sil’tronix Silicon Technologies, Archamps, France). Samples were prepared from a chloroform-methanol stock of 1 mg/ml DMPC:DMPG (1:1). Using Langmuir-Blodgett and Langmuir-Schaefer techniques, the two membrane leaflets of the bilayers were deposited sequentially. The surface pressure was kept at a constant 30 mN m^-1^ during the deposition, as described previously (Barker *et al.* 2016; Hubbard *et al.* 2017; Raasakka *et al.* 2017). All sample blocks were assembled into low-volume measurement flow cells, which were used for *in situ* exchange of solvent and injection of protein samples between reflectometric data collections (Junghans *et al.* 2015).

Neutron reflectometric measurements for wt-P0ct were performed as described (Raasakka *et al.* 2017). In short, the D17 neutron reflectometer at the Institut Laue-Langevin (Grenoble, France) was used for data collection at two incident angles (0.8° and 3.2°) (Cubitt and Fragneto 2002). All samples were kept at 30 °C with HBS buffer as the liquid phase, prepared at a final concentration of 95% (v/v) deuterium oxide (D_2_O, Sigma-Aldrich) and in H_2_O. The deposited bilayers were characterized, before and after the injection of P0ct, at three different solvent contrasts, varying the volume ratio of D_2_O and H_2_O in to the sample cell: (1) 95% D_2_O, (2) Si-matched water (SMW; 38% (v/v) D_2_O, 62% (v/v) H_2_O), and (3) 100% H_2_O. A 0.5 µM P0ct solution was allowed to interact with the membrane for 3 h whilst monitoring reflectivity, until no further changes were observed. Any excess P0ct was washed out from the bulk solution by exchanging 20 cell volumes of solvent slowly through the sample cell. Fitting was performed using Motofit in Igor Pro 7 (Nelson 2006).

The scattering length densities of the phospholipids were calculated from volume fractions of the lipid components obtained from molecular dynamics simulations (Armen *et al.* 1998), and for the proteins, they were calculated from the sequences and amino acid volumes (Zamyatnin 1972). The P0ct scattering length density, assuming 90% labile hydrogen exchange, was 3.227, 2.324, and 1.722 x 10^-6^ Å^-2^ in 95%, 38%, and 0% D_2_O, respectively. The errors in the structural parameters for each sublayer were derived from the maximum acceptable variation in the fitted thickness and lipid volume fraction that allowed a fit to be maintained, subject to a constant molecular area constraint required to maintain a planar bilayer geometry.

Details of the analysis of supported lipid membrane structure (Vacklin *et al.* 2005) and interaction with soluble proteins (Wacklin *et al.* 2016) using time-of-flight neutron reflection have been described previously. The fraction of P0 in the lipid bilayers was determined by a simultaneous fit to all contrasts, taking into account the change in protein scattering length density with solvent contrast due to H/D exchange of protons on polar residues with the solvent.

For mutant comparison to wt-P0ct, NR data for wt-P0ct and D224Y were collected on the INTER neutron reflectometer at ISIS Neutron and Muon Source (Didcot, United Kingdom) at two incident angles (0.7° and 2.3°) (Webster *et al.* 2006) covering a total q-range from 0.01 to 0.34 Å^−1^, with a resolution of Δq/q=0.03. The samples were prepared and handled as above.

## Acknowledgements

This work was financially supported by the Academy of Finland (Finland), the Jane and Aatos Erkko Foundation (Finland), and the Norwegian Research Council (SYNKNØYT program). This work has been supported by iNEXT, grant number 653706, funded by the Horizon 2020 programme of the European Commission. The research leading to this result has been supported by the project CALIPSOplus under the Grant Agreement 730872 from the EU Framework Programme for Research and Innovation HORIZON 2020. We gratefully acknowledge the synchrotron radiation facilities and the beamline support at ASTRID2 and EMBL/DESY, as well as the ILL (Proposal No. 8-02-745) and STFC/ISIS (Proposal No. 1620422; doi:10.5286/ISIS.E.95673822). We express our gratitude towards the Biocenter Oulu Proteomics and Protein Analysis Core Facility for providing access to mass spectrometric instrumentation, as well as the Biophysics, Structural Biology, and Screening (BiSS) facilities at the University of Bergen. Finally, we thank Anushik Safaryan for practical help with liposome preparation.

## Author contributions

Original text and figures: A.R., P.K.

Prepared mutant constructs: A.R., C.K.K., G.H.V., E.I.H.

Protein expression and purification: A.R., O.C.K.

Prepared samples and performed experiments: A.R., S.R., R.B., M.W.A.S., U.B.

Processed and analyzed data: A.R., R.B., U.B., H.W., N.J., S.V.H., P.K.

Study design: A.R., S.R., P.K.

Review & editing: A.R., S.R., R.B., H.W., N.J., S.V.H, P.K.

Supervision: P.K.

## Data availability

The datasets generated and analyzed during the current study are available from the corresponding author on reasonable request.

## Supplementary information

**Supplementary Fig. S1. Protein yield.** The purified protein amount from *E. coli* expression, shown as mg of protein obtained per 1 l of culture.

**Supplementary Fig. S2. The folding of P0ct variants in TFE, detergents, and poorly binding membrane compositions.** The folding of wt-P0ct and mutants was studied using SRCD spectropolarimetry in (a) 10% TFE, (b) 50% TFE, (c) 70% TFE, (d) 0.1% DPC, (e) 1% LDAO, (f) 1% OG, (g) DMPC, and (h) 9:1 DMPC:DMPG. The colour coding legend in panel (a) for each mutant trace also corresponds to all other traces in subsequent panels.

**Supplementary Fig. S3. P0ct variant-induced turbidimetry and diffraction.** (a) Turbidimetric analysis of 0.5 mM DMPC:DMPG (1:1) vesicles in the presence of 0 – 10 µM wt-P0ct and mutants. BSA was included as negative control. Error bars represent standard deviation. (b) Examples of Bragg peaks from the P0ct samples mixed with DMPC:DMPG (1:1) vesicles.

**Supplementary Fig. S4. EM analysis of P0ct D224Y.** Negatively stained samples of DMPC:DMPG (1:1) vesicles mixed with P0ct D224Y at (a) 1:100, (b) 1:200, and (c) 1:500 P/L ratios all display multilayered lipid structures.

**Supplementary Fig. S5. NR data of wt-P0ct and D224Y.** NR data for DMPC:DMPG (1:1)-bound wt-P0ct and D224Y. The data have been offset for clarity. Solvent contrasts are indicated for each trace on their right hand side. The D224Y H_2_O data is incomplete as reflectivity was collected at only one measurement angle (0.7°). The error bars denote standard deviation.

**Supplementary Table 1.** DLS parameters.

| Protein variant | wt-P0ct | T216ER | A221T | D224Y | R227S | K236E | K236del |
| --- | --- | --- | --- | --- | --- | --- | --- |
| Sample buffer* | HBS | HBS | HBS | TBS | HBS | HBS | HBS |
| $R_h$ (nm) | 2.96 | 2.87 | 2.72 | 2.26 | 2.93 | 2.88 | 2.80 |

\*HBS, 20 mM HEPES, 150 mM NaCl, pH 7.5; TBS, 20 mM Tris-HCl, 300 mM NaCl, pH 8.5.

**Supplementary Table 2.** SAXS parameters.

| Data collection parameters |  |  |  |  |  |  |  |
| --- | --- | --- | --- | --- | --- | --- | --- |
| Instrument | P12, PETRAIII, DESY |  |  |  |  |  |  |
| Wavelength (nm) | 0.124 |  |  |  |  |  |  |
| Angular range (nm<sup>-1</sup>) | 0.0403 - 7.3195 |  |  |  |  |  |  |
| Exposure time (s) | 0.045 |  |  |  |  |  |  |
| Measurement temperature (°C) | 10 |  |  |  |  |  |  |
| Protein variant | wt-P0ct | T216ER | A221T | D224Y | R227S | K236E | K236del |
| Concentration range (mg ml<sup>-1</sup>) | 0.3 - 1.2 | 1.0 - 3.8 | 2.0 - 8.0 | 0.5 - 2.1 | 1.7 - 6.8 | 3.5 - 12.9 | 2.3 - 9.3 |
| Sample buffer* | HBS | HBS | HBS | TBS | HBS | HBS | HBS |
| Structural parameters |  |  |  |  |  |  |  |
| $I_0$ (relative) [from p(r)] | 58.90 | 64.72 | 59.06 | 58.27 | 56.14 | 62.73 | 58.17 |
| $R_g$ (nm) [from p(r)] | 2.57 | 2.50 | 2.40 | 2.73 | 2.42 | 2.41 | 2.41 |
| $I_0$ (relative) [from Guinier] | 58.25 | 63.87 | 58.35 | 57.38 | 55.21 | 61.61 | 57.30 |
| $R_g$ (nm) [from Guinier] | 2.39 | 2.33 | 2.26 | 2.43 | 2.25 | 2.23 | 2.23 |
| $D_{max}$ (nm) [from GNOM] | 9.59 | 9.21 | 9.59 | 11.57 | 8.96 | 10.34 | 10.69 |
| Molecular mass determination |  |  |  |  |  |  |  |
| Molecular mass $M_r$ (kDa) [from $I_0$ using p(r)] | 7.87 | 8.65 | 7.89 | 7.79 | 7.50 | 8.38 | 7.78 |
| Molecular mass $M_r$ (kDa) [from $I_0$ using Guinier] | 7.79 | 8.54 | 7.80 | 7.67 | 7.38 | 8.24 | 7.66 |
| Theoretical $M_r$ from sequence (kDa) | 7.99 | 8.17 | 8.02 | 8.04 | 7.92 | 7.99 | 7.86 |
| Software |  |  |  |  |  |  |  |
| Primary data reduction & processing | PRIMUS |  |  |  |  |  |  |

\*HBS, 20 mM HEPES, 150 mM NaCl, pH 7.5; TBS, 20 mM Tris-HCl, 300 mM NaCl, pH 8.5.

