## Supplementary figures and images for "Neuropathy-related mutations alter the membrane binding properties of the human myelin protein P0 cytoplasmic tail"

### Supplementary Figure 1

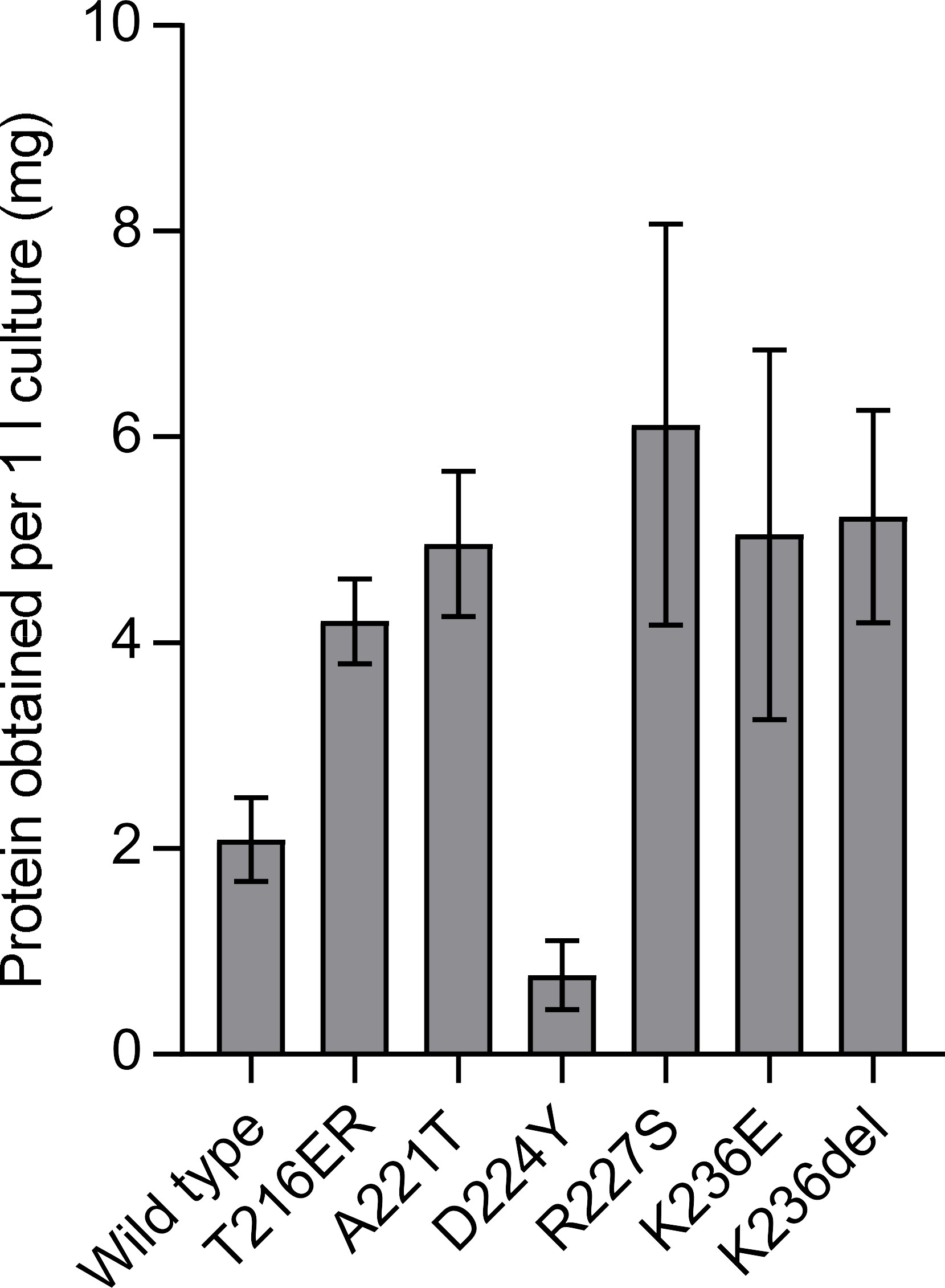

### Supplementary Figure 2

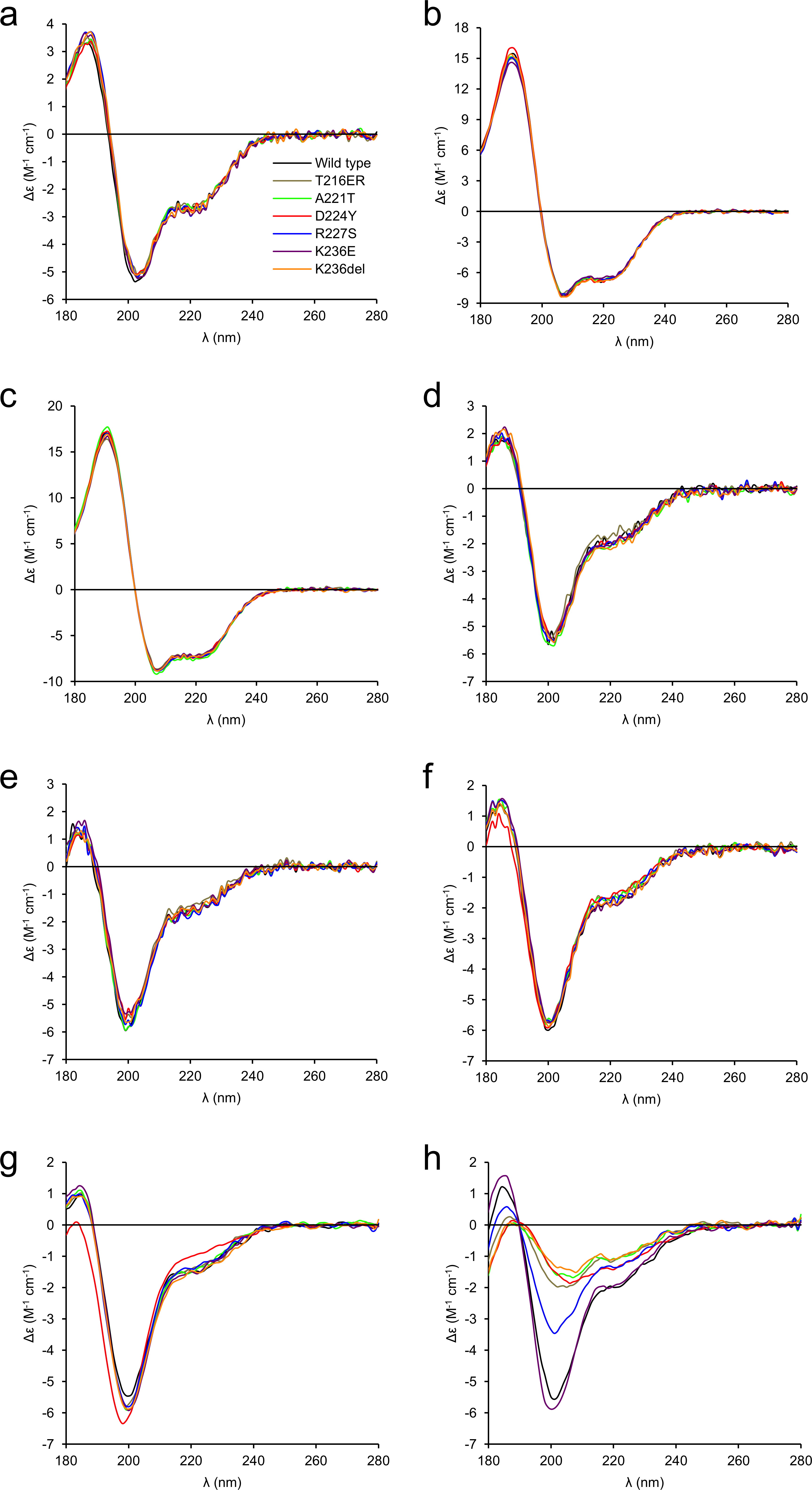

### Supplementary Figure 3

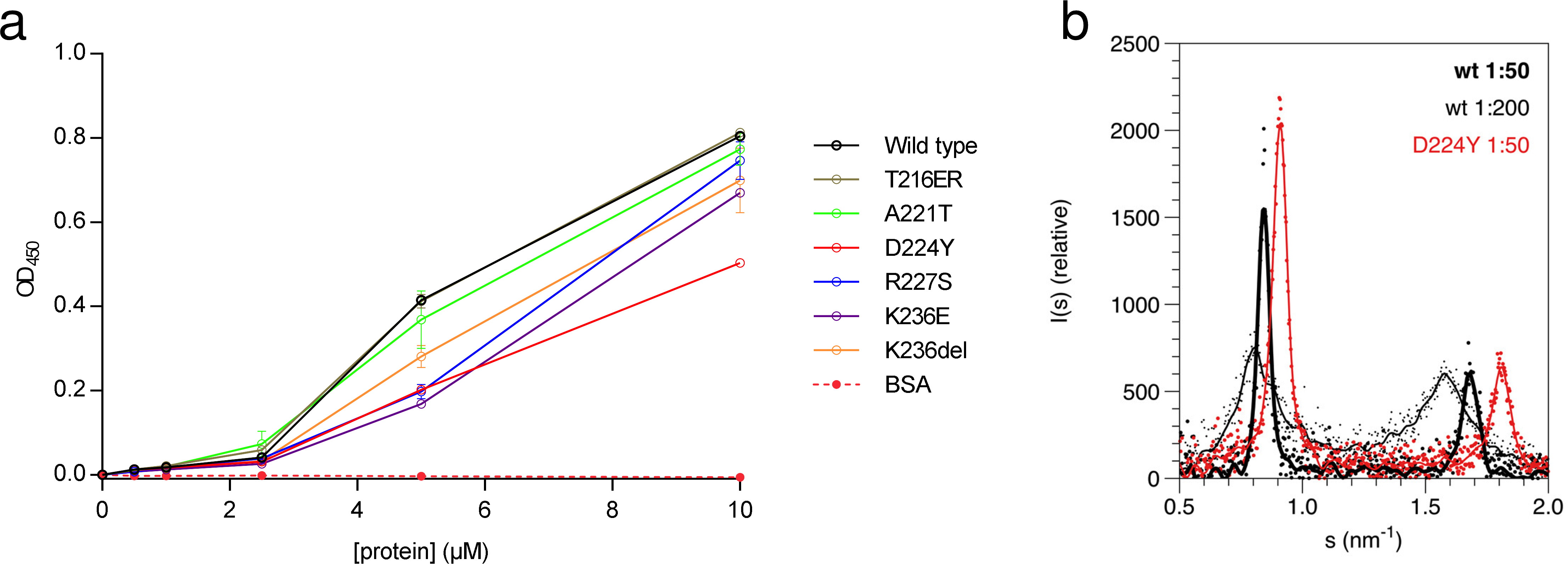

### Supplementary Figure 4

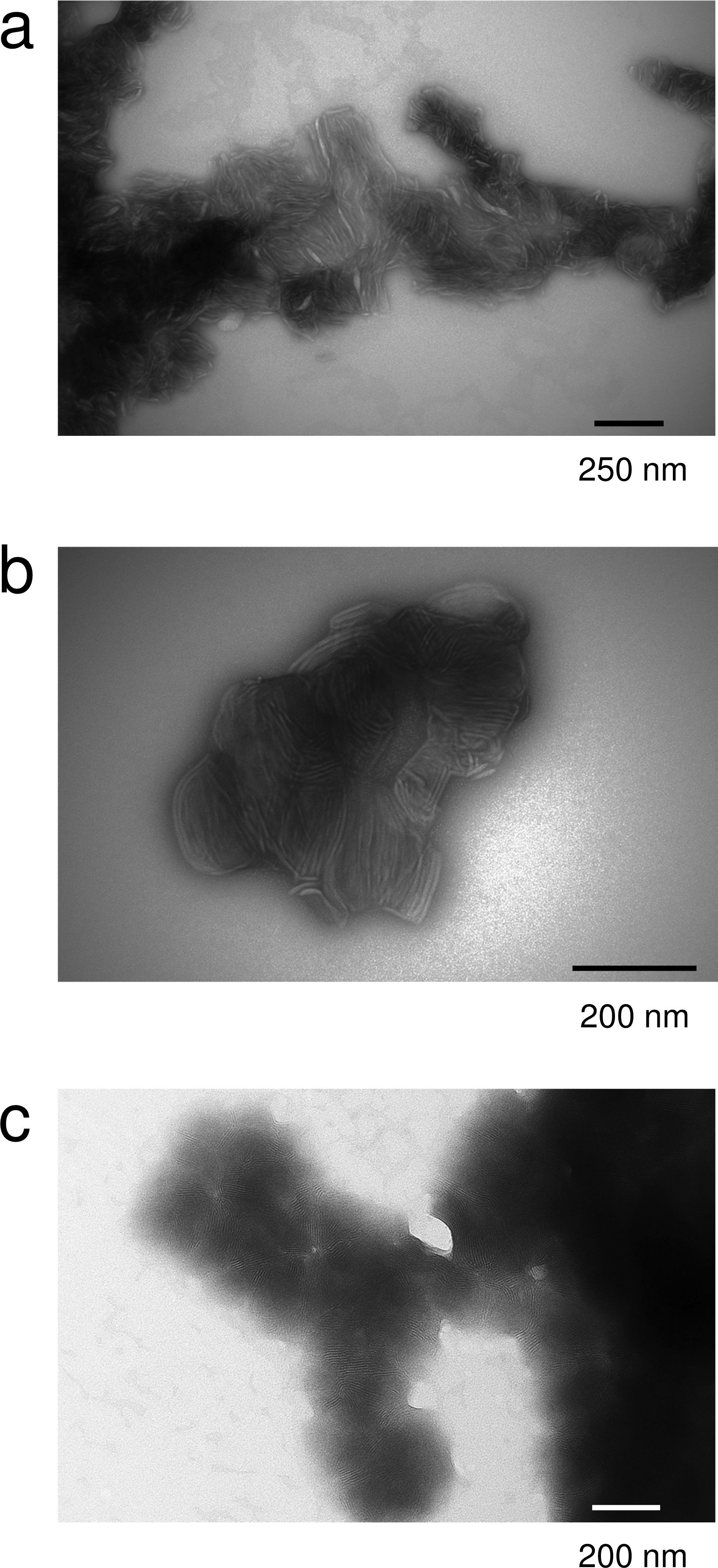

### Supplementary Figure 5

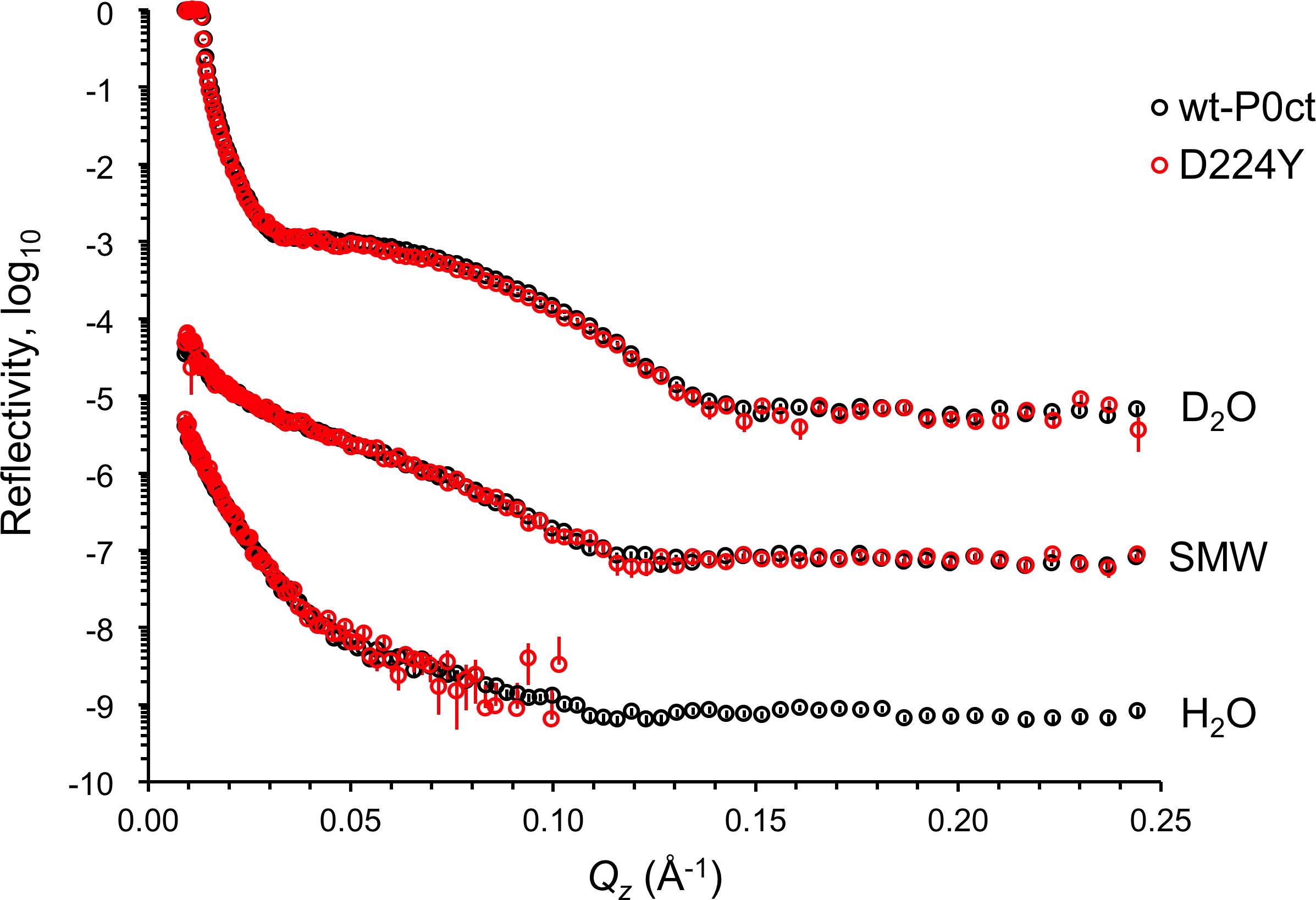
